# Courtship ritual of male and female nuclei during fertilisation in *Neurospora crassa*

**DOI:** 10.1101/2021.03.10.434724

**Authors:** Sylvain Brun, Hsiao-Che Kuo, Chris E. Jeffree, Darren D. Thomson, Nick Read

## Abstract

Sexual reproduction is a key process influencing the evolution and adaptation of animals, plants and many eukaryotic microorganisms, such as fungi. Mycologists have described the different fungal fruiting bodies, while geneticists have partly unravelled the regulation of sexual development. However, the sequential fungal cell biology of fertilisation and the associated nuclear dynamics after plasmogamy are poorly understood. Using histone-fluorescent parental isolates, we tracked male and female nuclei during fertilisation in the model ascomycetes *Neurospora crassa* using live-cell-imaging. This study unravels the behaviour of trichogyne resident female nuclei and the extraordinary manner that male nuclei migrate up the trichogyne to the protoperithecium. Our observations raise new fundamental questions about the *modus operandi* of nuclei movements during sexual reproduction, male and female nuclear identity, guidance of nuclei within the trichogyne and, unexpectedly, the avoidance of “polyspermy” in fungi. The spatio-temporal dynamics of male nuclei within the trichogyne following plasmogamy are also described, where the speed and the deformation of male nuclei are of the most dramatic observed to date in a living organism.

## Introduction

Fungi are living microorganisms that considerably impact human life. These adaptive eukaryotes have been shown to thrive in the damaged Chernobyl radioactive reactors (Dighton et al., 2008), space (Onofri et al., 2015) and extreme temperatures (Paulussen et al., 2017). Their impressive growth capacity is used for biotechnology and industrial purposes (Cairns et al., 2018; Chambergo and Valencia, 2016). Fungal plant pathogens are responsible for the loss of billions of dollars in food security due to crop losses worldwide, while human fungal pathogens are a major health problem that threatens the lives of millions (Bongomin et al., 2017; Brown et al., 2012; Fisher et al., 2012; Köhler et al., 2017). However, our knowledge of whole reproductive life cycles in the fungal kingdom is only partial. Sexual reproduction is a key event in the fungal life cycle, where evolution and selection rely on its creation of novel and beneficial genetic combinations in the organism (Heitman, 2010; Heitman et al., 2013). The fungal reproductive cycle relies on elaborate genetic regulatory systems as well as complex cytological events (Bennett and Turgeon, 2016; Debuchy et al., 2010; Ni et al., 2011). Understanding reproduction provides means to mitigate fungal threats and enhance bio-technological processes.

In heterothallic model organisms such as the filamentous fungus *Neurospora crassa* and *Saccharomyces cerevisiae* yeast, sexual reproduction only occurs with opposite mating type partners (*mat a* and *mat A* in *N. crassa*) (Dodge, 1927). In *N. crassa*, Dodge identified conidia as the male partner and the ascogonium as the female partner (Dodge, 1935). The ascogonium is embedded within a multi-hyphal female structure called the protoperithecium (Lichius et al., 2012). The ascogonium produces specialised hyphae called trichogynes to facilitate fertilisation. Trichogynes are chemotropically attracted by a diffusible chemical signal emitted by conidia (Bistis, 1981, 1983). The trichogynes eventually fuse to their macro- or micro-conidial mate, allowing successful fertilisation and development of the protoperithecium into a perithecium (Backus, 1939).

Trichogynes express G Protein-Coupled Receptors (GPCRs) at their surface which respond to the pheromone signal emitted by conidia (Bobrowicz et al., 2002; Jones and Bennett, 2011; Kim and Borkovich, 2004, 2006; Kim et al., 2002, 2012; Krystofova and Borkovich, 2006). In ascomycetes, the expression of both the pheromone and the GPCRs are under the control of the *mat* locus (Debuchy et al., 2010). Trichogynes, which are attracted by the pheromone signal bind to, coil around and eventually fuse with male conidia, eventually establishing plasmogamy. Although the migration of male nuclei has never been observed, it is assumed that once within the female trichogyne, male nuclei (of the opposite sex mating type) migrate up the trichogyne to the ascogonium where they proliferate together with female nuclei eventually forming the dikaryotic bag (Raju, 1992). Karyogamy and meiosis take place once two nuclei of each mating type exit the dikaryotic bag to enter the ascogenous cell. Although fertilisation and development of perithecia have been imaged in several ascomycete models, merely by electron micrographs (Backus, 1939; Dodge, 1935; Harris et al., 1975; Lord and Read, 2011; Mai, 1976, 1976; Read, 1983; Read and Beckett, 1996), the actual behaviour of male and female nuclei in this fundamental process has never been tracked in living cells.

In the lab, fertilisation in *N. crassa* can be observed a few hours after inoculating the protoperithecia with a suspension of opposite mating-type conidia, making this fungus a well-suited experimental model to study fertilisation (Backus, 1939; Bistis, 1981, 1983). Here, using parental *N. crassa* strains in which nuclei were visualised with either the synthetic Green Fluorescent Protein (H1-sGFP) (Freitag et al., 2004) or the tdimer Red Fluorescent Protein (H1-RFP) fused to the histone H1 protein, we achieved for the first time live-cell imaging of male and female nuclei during fertilisation. Our observations highlight remarkable differences in the behaviour of nuclei from different parental origins within the same cellular cytoplasm. Moreover, we imaged the astounding dynamism of male nuclei as they migrate by repeated stretching and contraction up the trichogyne passing by immobilised female nuclei. Finally, our results raise the question of the signalling network controlling the movements of both male and female nuclei throughout fertilisation. In particular, we provide evidence for a system preventing the protoperithecium to be fertilized by several male nuclei, which operates similarly to systems avoiding polyspermy in mammals (Firman, 2018).

## Results

### Chemotropic growth of trichogynes

To visualise the fertilisation process in *N. crassa*, we adapted existing experimental settings (Bistis, 1981). This involved inoculating 1.5 x 1.5 cm agar blocks bearing 5-7 day-old protoperithecia producing “female” trichogynes with a mixed suspension of “male” macro- and microconidia from the opposite mating type. After 6h inoculation, the agar blocks were inverted onto a micro-slide chamber containing a droplet of Vogel’s medium and imaged on either an inverted Nikon TE-2000 widefield or Leica SP8x confocal microscope. Crosses were performed in both directions: ♀ *mat A* X ♂ *mat a* as well as ♀ *mat a* X ♂ *mat A*. We observed no differences between the two cross types. Trichogynes were identified via the following morphological features (Movies S1 & S2): reaching several hundred micrometres (Figures 1, 2, 4A, 5A and 6); exhibiting tropic and sinusoidal growth towards conidia (Figures 1B, 1C, 2B, Movies S1 & S2), at a reduced extension rate of 1.1 ± 0.2 µm / min, compared to vegetative hyphae at the colony edge (67 µm / min) (Ryan et al., 1943); and exhibited coiled growth around conidia. Both micro- and macro-conidia were found to attract and fuse with trichogynes. Hence, the coiling of the trichogyne around the conidium turned out to be a hallmark characteristic of “successful” fertilisation (Figures 1C, 4A and 6).

**Figure 1:**
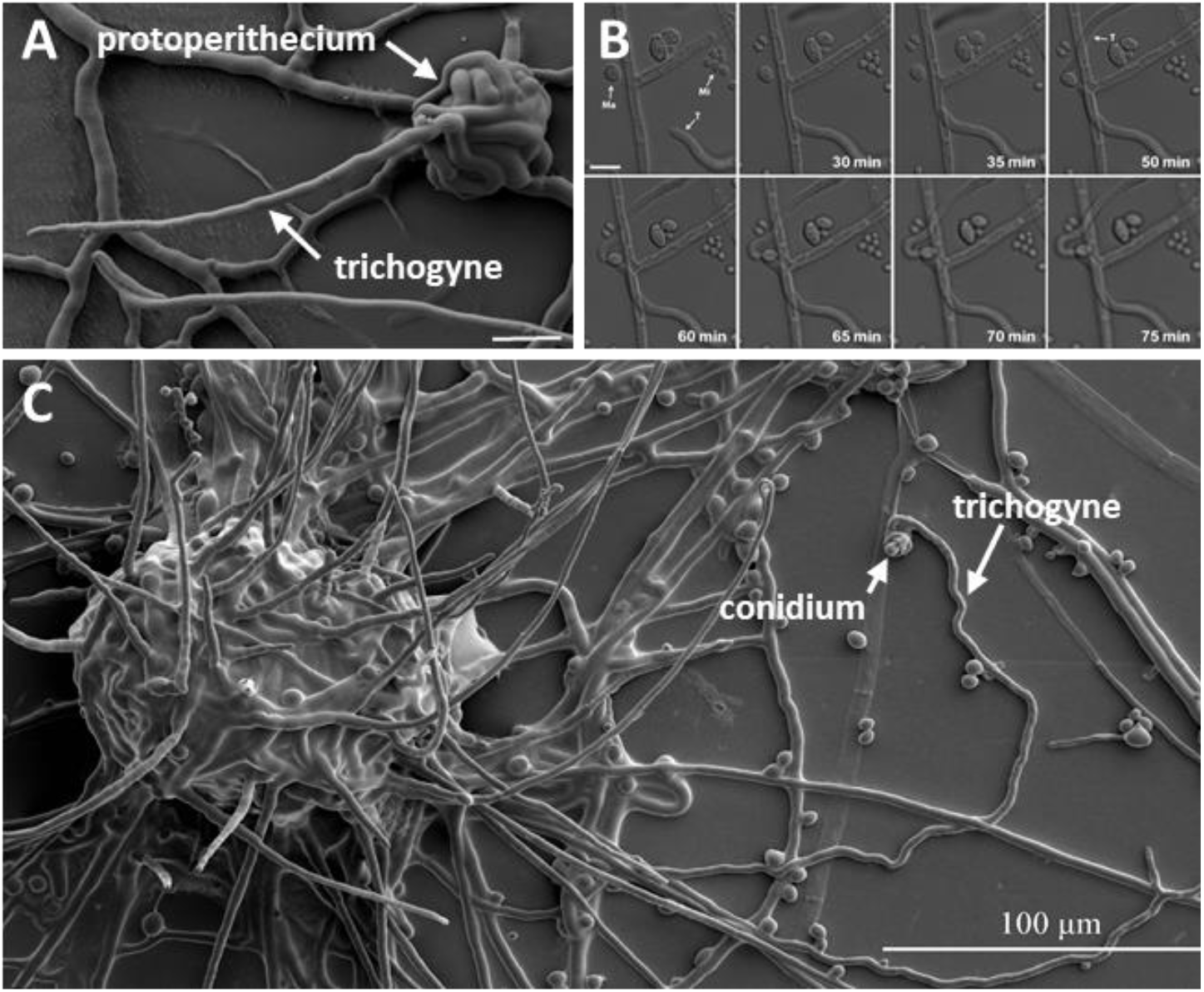
Trichogynes and chemotropic growth in *N. crassa*. **(A)** A SEM image of a young (3 days old) protoperithecium emitting a single trichogyne; scale bar 10 µm. **(B)** Time-lapse sequence of the tropic growth of two trichogynes (T) toward one isolated macroconidium (Ma) while ignoring other macroconidia and microconidia (Mi). The trichogynes are not attracted towards the microconidia in this field of view. Scale bar 10 µm. **(C)** SEM image of a 5-7 days old protoperithecium, where one trichogyne has been attracted and coiled around a conidium; scale bar 100 µm.

Since conidia are randomly spread on trichogyne-bearing agar blocks, the vicinity of protoperithecia to individual conidia or groups of conidia varied. As a result, trichogynes from individual protoperithecia were observed to extend and fuse with conidia located between a few microns to half a millimeter from their site of origin. We observed very long trichogynes (> 500 µm) from a protoperithecium fusing with a conidium located close to another protoperithecium. As observed in Figure 1B, trichogynes do not necessarily fuse with the first conidia encountered but can instead be attracted to other conidia further away. Interestingly, an isolated singular macroconidium, in vicinity to clusters of macroconidia and microconidia, attracted two trichogynes, suggesting heterogeneity in the conidial ability to attract trichogynes (Figure 1B).

### Behaviour of female nuclei in the course of fertilisation

We imaged *H1-sGFP* tagged female nuclei i) during the tropic growth of trichogynes towards male conidia, ii) after the trichogyne has coiled the conidium and received the first *mat a H1-RFP* male nucleus, and iii) when male nuclei migrate up to the trichogyne base. The first contact between conidia and trichogynes were regularly observed 4-6 h after inoculation with the conidial suspension. Entry of male nuclei into trichogynes and protoperithecia occurred 6-8 h and 8-10 h after inoculation, respectively.

Importantly, female and male H1 histone proteins, fused with either green (syntheticGFP; sGFP) or red (tdimerRed; RFP) fluorescent proteins, never mixed in the same nuclei. This enabled us to exclusively follow the GFP-female and RFP-male nuclei throughout the fertilisation process in our imaging experiments. Female nuclei were located and aligned throughout the trichogyne axis (Figure 2 and Movie S2). During tropic trichogyne growth, GFP female nuclei displayed an overall anterograde movement at rates similar to the extension rate of the trichogyne. From Figure 2 & Movie S2, upon trichogyne-conidium contact (t = 36 min), it took 23 min for the trichogyne to complete conidium-coiling and cease growth (t = 59 min). Next, the first *H1-RFP* conidial nucleus took 53 min to enter into the trichogyne (t =1 h 52 min), followed by a second one 10 min later (t = 2 h 02 min), indicating that plasmogamy had occurred. Upon cessation of trichogyne growth at its targeted conidium, the anterograde movement of female nuclei switched to an oscillatory motion and then became visually immobilised 4 min before the entry of the first male nucleus (t= 1 h 48 min). Nuclei speed measurements revealed that female nuclei movements did not fully stop but were reduced. For sake of simplicity, we will refer to this phase as female nuclei immobilisation. These female nuclear dynamics suggest stepwise signalling events anticipating the entry of the male nucleus. Female nuclei remained immobilised while male nuclei migrated up the trichogyne. Finally, 56 min after the first male RFP nucleus entered the trichogyne all immobilised female GFP nuclei synchronously began moving backward (retrograde movement) up the trichogyne (t = 2 h 44 min). These female nuclear dynamics were consistent with every fertilisation event analysed (n >10). We quantified female nuclei mobility before and after entry of male nuclei into trichogynes for at least one fertilisation event. By tracking nuclei in Movies S3a (before entry) & S3b (after entry), we measured the speed of GFP female nuclei within trichogynes after trichogyne growth had ceased at the conidium and after female nuclei were immobilised. These dynamics were compared to the mobility of female nuclei located in surrounding hyphae, likely vegetative ones since initial nuclei behaviour in these hyphae were different (Figure 3, Movie S3a & b).

**Figure 2:**
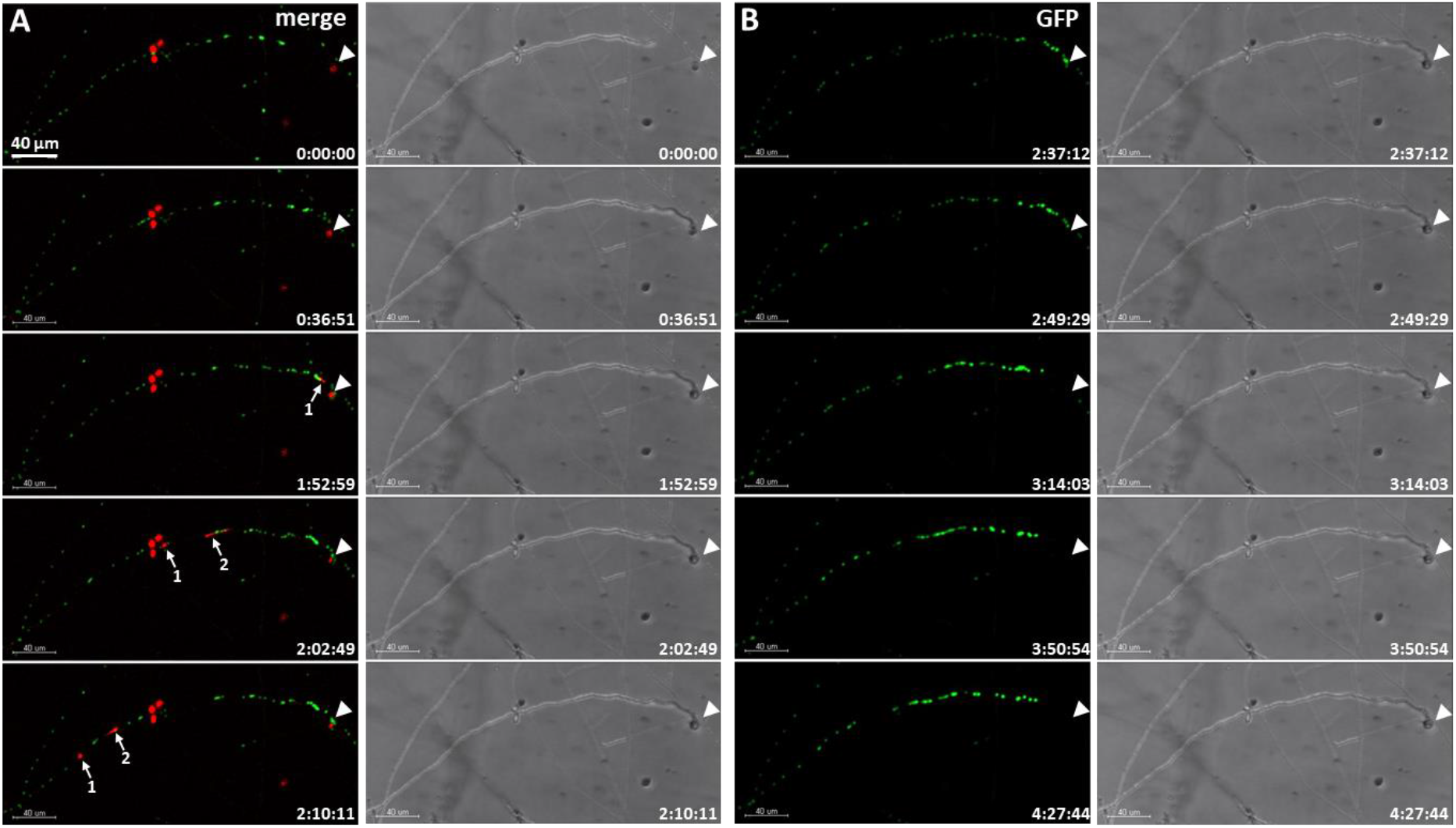
Time lapse imaging of male and female nuclei before and after plasmogamy (Movie S2). Female *mat A H1-sGFP* was fertilized with *mat a H1-RFP* male conidia. A) merged fluorescence and brightfield confocal images (maximum intensity projected Z-stacks) during the initial stages of fertilisation. The female trichogyne bearing sGFP nuclei trophically grows to one macroconidium containing RFP nuclei (arrowhead). Two RFP male nuclei from the macroconidium (arrows 1 and 2) enter and migrate up the female trichogyne. B) GFP fluorescence and brightfield confocal images (maximum intensity projected z-stacks) during latter stages of fertilisation after RFP male nuclei have left the field of view. Female sGFP nuclei undergo retrograde movement up the trichogyne from its coiled tip (arrowhead). Scale bar = 40 µm.

The average speed of hyphal female nuclei was significantly higher (2.04 ± 2.36 µm / min) compared to trichogyne female nuclei (1.03 ± 1.00 µm / min; Figure 3 p<10^−3^), prior to male nucleus entry. Strikingly, the speed of both hyphal and trichogyne female nuclei significantly decreased, after male nucleus entry, to 1.33 ± 1.07 µm / min (p<10^−3^) and 0.82 ± 0.86 µm / min (p<10^−3^), respectively. These data confirmed the immobilisation of female nuclei observed immediately prior entry of male nuclei into the trichogyne.

**Figure 3:**
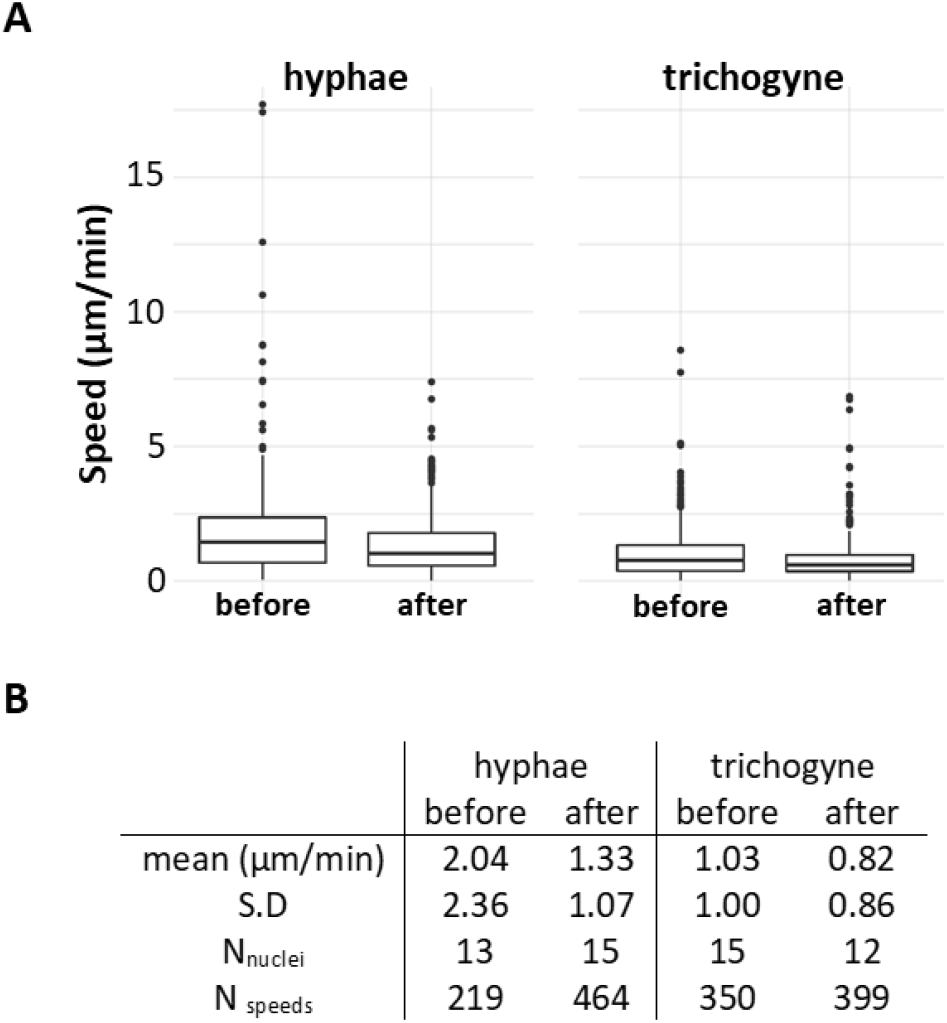
Female nuclear speeds before and after male nucleus entry into the trichogyne. Analysis of the time lapse Movies S3a & b.(A) Plot, (B) numerical values. The speed of the green-labelled *H1-sGFP* female nuclei of the trichogyne as well as of the vegetative hyphae in the same field of view were measured at every time frames before (Movie S3a) and after (Movie S3b) entry of male nucleus into the trichogyne. S.D, Standard Deviation. N_nuclei_, number of analysed nuclei in the field of view. N_speed_, number of nuclei speeds measured. Results from the ANOVA (on log values) indicate that the effect of target site is significant (P < 0.001), with larger values in hyphae (1.56 ± 1.63, n = 683) compared to trichogyne (0.92 ± 0.94, n = 749); likewise, the effect of time of observation is significant (P < 0.001) with larger values before (1.42 ± 1.73, n = 569) compared to after (1.10 ± 1.01, n = 863); these two effects are independent one from the other as confirmed by the non-significant interaction (P = 0.431). Pairwise t-tests with pooled SD indicate that all pairs of means were different at the 5% level.

The shape of female *H1-sGFP* fluorescent nuclei in both the analysed hyphae and in the trichogyne changed from ovoid to more-rounded during their immobilised phase, as observed in Movies S3a & S3b and reverted to ovoid at the resumption of movement (end of Movie S5). During this latter phase (Figure 5; Movie S5), the recovered mobility in female nuclei followed by their retrograde migration up the trichogyne was exemplified by the behaviour of the nucleus, close to the conidium in Figure 5 and Movie S5 (the top one). This exemplar nucleus became pear-shaped while being pulled in the retrograde direction. These data describe for the first time the dynamic nature of resident female nuclei within trichogynes during sexual fertilisation in *N*.*crassa*. Moreover, the signal that triggers female nuclear immobilisation is not restricted to the trichogyne subjected to plasmogamy, suggesting a possible diffusible signal, which can pause the nuclear dynamics of this filamentous fungus.

### Nuclear morphology and movement of residual conidial male nuclei

Using the very same protocol described previously, we aimed to image *mat a H1-RFP* male nuclei at the onset of their entry into female *mat A H1-sGFP* trichogynes. Four to six hours after inoculation with conidia of opposite mating type, scarce trichogynes coiling around conidia (fertilisation events) were observed on inoculated agar blocks. In order to image early steps of male nuclei migration, image acquisitions were started as soon as coil fertilisation hallmarks were found. This way, we observed that male RFP nuclei were highly mobile, rotating while still enclosed within the conidium (Figure 4 & 5, Movies S4 & S5). Eventually, these rotating movements accelerated until a nucleus (sometimes one of several within the macroconidium) started to repetitively elongate towards the entry of the trichogyne. In the particular case of the conidium imaged in Figure 4 (Movie S4), no male nucleus entered the trichogyne. Same observation was made for the nucleus remaining in the conidium in Figure 5 (Movie S5). We suspected that lack of male nucleus migration could be due to photo-toxicity and that real-time acquisition had to be limited in time before the beginning of male nuclei migration.

**Figure 4:**
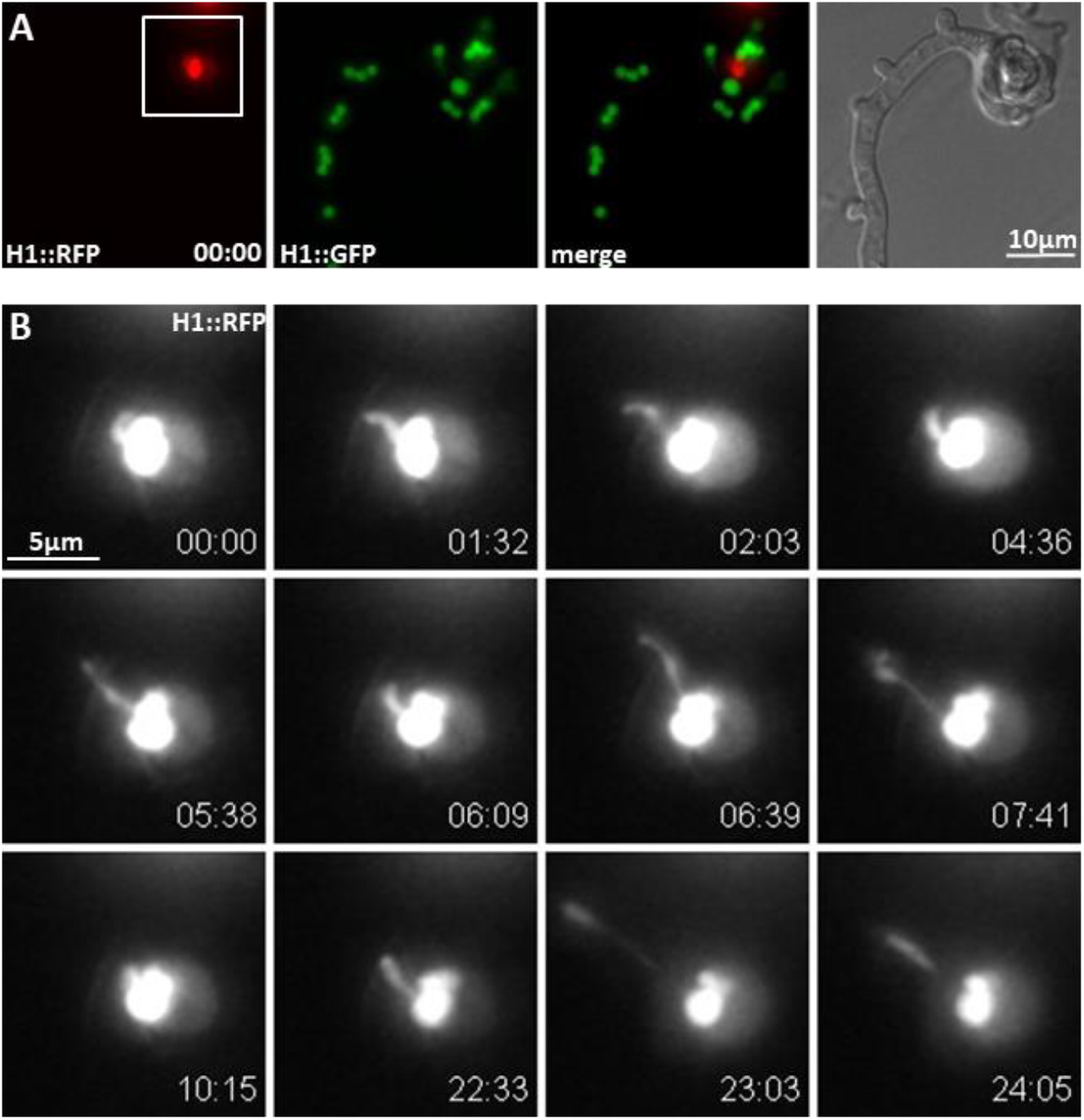
Time course of male nuclear mobility within the conidium upon fertilisation (Movie S4). **(A)** Widefield fluorescence and brightfield images of *H1-sGFP* female trichogyne fertilisation by a *H1-RFP* conidium. **(B)** Higher magnification of the residual RFP conidial nuclei (scare box in A) after the first nucleus entered and migrated up the trichogyne (not shown). The imaged residual nucleus stretches and moves around within the conidium but failed to migrate up the trichogyne. Time scale: min:sec. Scale bars = 10 µm in A and 5 µm in B.

### The inch-worm movement of male nuclei

To minimise photo-toxicity to the fertilisation process, fertilisation events were checked infrequently in order to preserve the integrity of subsequent live-cell imaging of the male nuclear migration. This extra care allowed observation of several male nuclei migration events into trichogynes. The trichogyne entry process was not straightforward, male nuclei do not directly enter the trichogyne, but instead paused frequently prior to entry. It was frequently observed that individual male nuclei stretched and retracted in and out of the trichogyne several times without leaving the conidium (data not shown). This “inch-worm” nuclear movement is illustrated in Figure 5 and Movie S5, where the RFP male nucleus moved in an “inch-worm” manner, repetitively stretching and contracting while migrating up the trichogyne. Stretching was so dramatic that male nuclei seemed sometimes split into two (Figure 5; from 3 min 18 sec - 4 min 03 sec; Movie S5). The way male nuclei stretched suggested that the front is submitted to pulling mechanical strain during migration (see discussion). Nuclear elongation was especially striking when the male nucleus passed through septa. In Figure 5, the male nucleus passed through the first septum (Figure 5; t=2 min 10 sec to t=4 min 03 sec) and remarkably stretched to a maximum length of 40 µm (t= 3 min 27 sec; Figure 5), where its apical end was pulled while its rear end was “stuck” at the septal pore. To understand this process more, we measured the speed of each nuclear apex (front and rear) in the movie. The maximum speed measured for the front side was 85 µm / min while the maximum speed of the rear side reached 130 µm / min (Figure 5B). These data highlighted the remarkable spatio-temporal dynamics and nature of migrating fungal nuclei during fertilisation in *N. crassa*.

**Figure 5:**
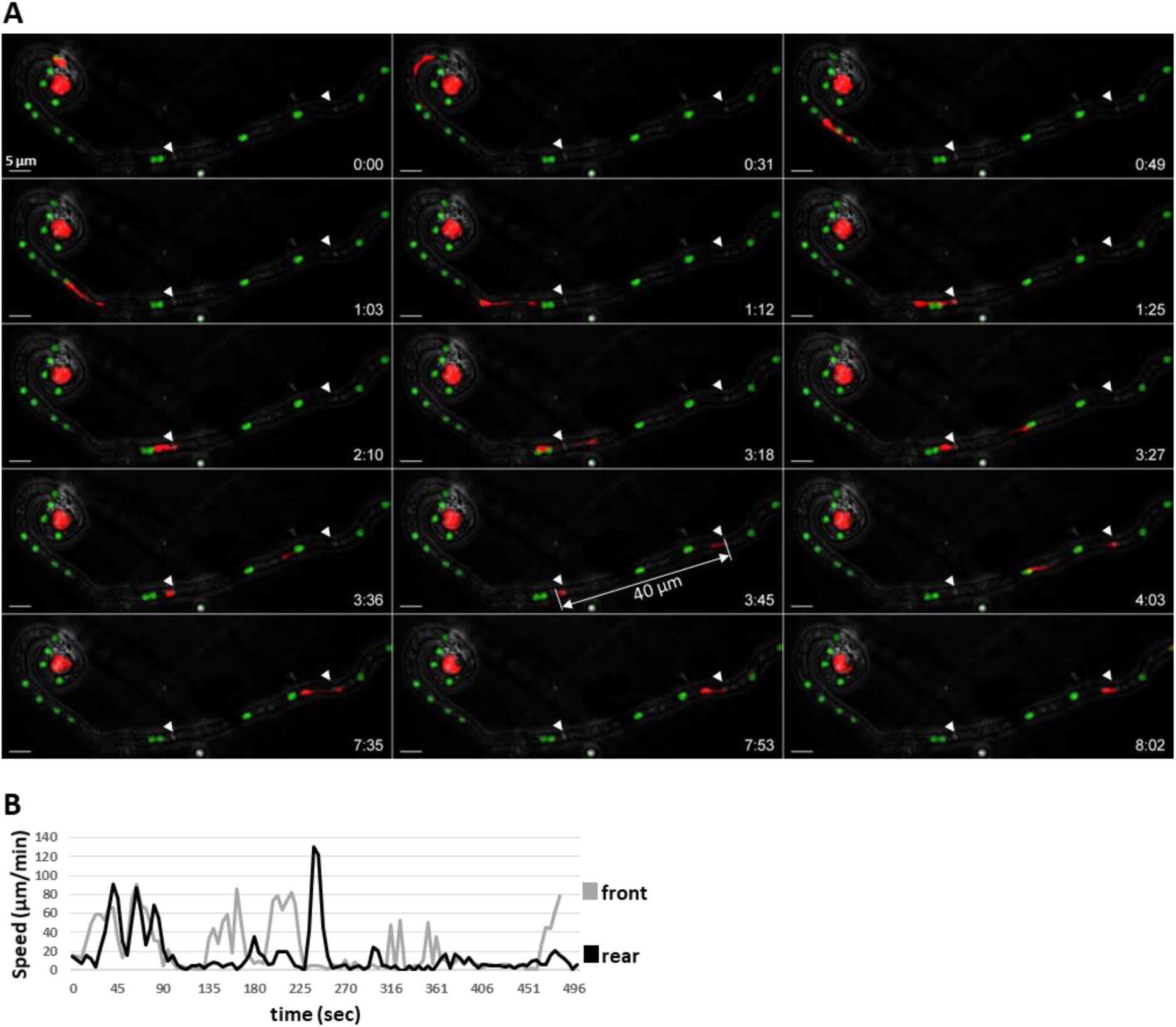
Time course of male nucleus movement across the trichogyne (Movie S5). **(A)** Merged confocal images (maximum intensity projected z-stacks) of the fertilisation of a *mat A H1-sGFP* female trichogyne by *mat a H1-RFP* conidium. The RFP nucleus enters and migrates up the female trichogyne successively, stretching and contracting in an inch-worm like manner. The remarkable stretching (40 µm) of the male nucleus crossing the first septa (arrowheads) is indicated. Scale bar = 5 µm. **(B)** Velocity analysis of the exemplar stretching RFP nucleus at the front (grey) and rear points (black).

### Fertilisation of the protoperithecium by male nuclei

After trichogyne entry, the conidial nuclei travel to the protoperithecium where the next steps of sexual reproduction such as karyogamy, meiosis and ascosporogenesis take place. In *N. crassa*, macroconidia are typically multi-nucleate. We reproducibly observed that fertilisation by macroconidia involved discharge of several nuclei (up to three nuclei per conidium in Figure 6) from a single conidium into a trichogyne (Figure 6 & Movie S6). Furthermore, we observed that trichogynes can branch and that those branches are capable of binding, coiling and fusing with multiple conidia, at least two (A and B; Figure 6 & Movie S6). In the trichogyne of Figure 6, male nuclei were tracked from the conidia to the protoperithecium over a distance of up to 574 µm (Figure 6; Conidia A). Three male RFP nuclei entered the trichogyne from conidium A in 35 min (Figure 6; labelled 1, 2 and 3; from t = 46 min to t = 1 h 21 min) followed by three additional nuclei from conidium B which were released in only 6 min (Figure 6; labelled 4, 5 and 6; from t = 1 h 25 min to t = 1 h 31 min). Eventually, the presence of RFP nucleus 4 was detected within the protoperithecium (Figure 6; t= 1h 53 min). We assumed the nucleus entering the protoperithecium was nucleus 4 based on its proximity and the fact that we never observed “overtaking” behaviour in any male nucleus in our experiments. Strikingly, although the remaining male nuclei continued their migration up the trichogyne, all of them stalled and accumulated upstream of the now-fertilized perithecium (Figure 6; t= 4hr 24min; Movie S6). It was also the case in Movie S7 where two conidial male nuclei migrate up the trichogyne and only one entered into the protoperithecium (Movie S7). Accordingly, it was frequent to observe immobile RFP male nuclei in trichogynes when observed late in the experiment (between 10 and 14 hours) presumably after protoprithecia were fertilised (data not shown). These data led us to assume that entry of a first male nucleus into a protoperithecium inhibited the entry of the others. A second important feature provoked by entry of male nuclei into prothoperithecia was the quantified increase of volume of the latter (Figure 6; from t = 1 h 53 min; Figure 7). Finally, we detected a second RFP foci signal in the core of the protoperithecium twice (Figure 6; t = 2 h 25 min - 4h 27 min; Movie S7; 4 h 22 min 52 sec). Careful analysis of the images did not indicate any entry of any of the remaining nuclei. Thus, appearance of this second RFP foci signal 32 min after initial protoperithecium entry in Figure 6 (Movie S6) may indicate that this male nucleus 4 has divided. However, we cannot exclude that appearance of a second foci within the protoperithecium may be due to the stretching of the nucleus. Altogether, these data suggested that entry of male nuclei into protoperithecia triggered at least two signals: a first one in order to avoid polyspermy by blocking multiple fertilisation of the protoperithecium and a second signal committing the protoperithecium into a growing and developing perithecium.

**Figure 6:**
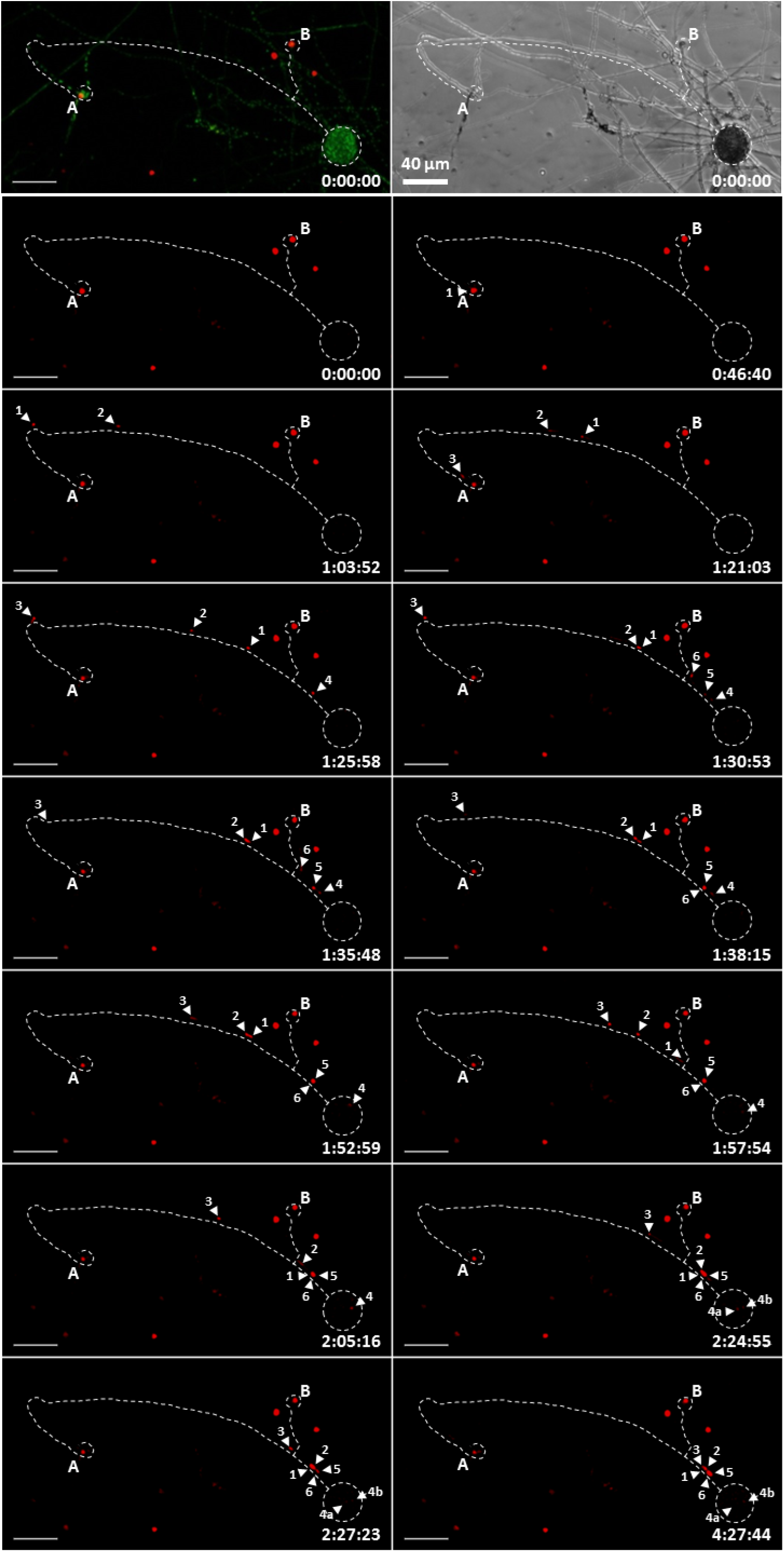
Time course of the fertilisation of a protoperithecium by a single male nucleus (Movie S6). Confocal microscopy of the fertilisation of H1-sGFP female with H1-RFP conidial suspension. First row shows merged GFP female nuclei and RFP male nuclei with a brightfield representation at t = 0 min. Subsequent timelapse RFP images contain a dotted trace of the trichogyne and protoperithecium. Two conidia (A and B) have attracted and fused with a branched trichogyne emitted by the protoperithecium at the bottom-right corner. Three nuclei (1, 2 and 3) from conidium A and three nuclei (4, 5 and 6) from conidium B successively enter the trichogyne. Nucleus number 4 enters the protoperithecium core and eventually divides (4a and 4b). Scale bar = 40 µm; time scale = h:min:sec.

**Figure 7:**
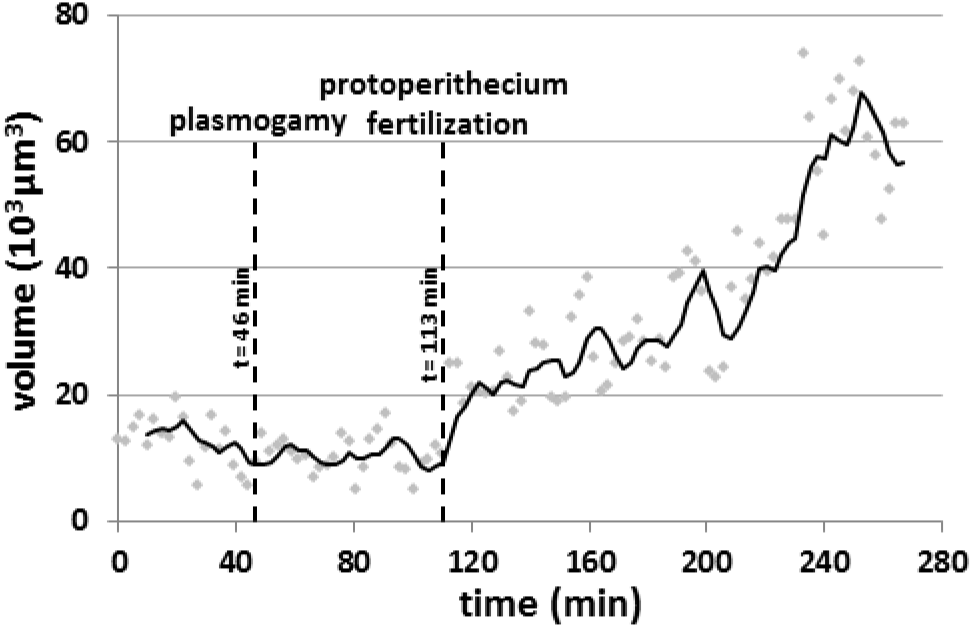
volume increase of protoperithecium after its fertilisation by entry of one male nucleus. Analysis of segmented images of *H1-sGFP* fertilisation by *H1-RFP* conidial suspension in Movie S6 (Figure 6). The green signal of the *H1-sGFP* nuclei composing the protoperithecium has been segmented in order to evaluate the volume of the perithecium in the course of fertilisation. The time of entry of the first *H1-RFP* male nucleus into the trichogyne (plasmogamy) as well as the time of entry of the nucleus number 4 into the protoperithecium (protoperithecium fertilisation) are indicated. Dots individual data points and black line = moving average.

### The orienteering race of male nuclei up the trichogyne

Another astonishing behaviour of conidial male nuclei was their capacity to rapidly switch between anterograde and retrograde movement during trichogyne migration (Movie S8 and S9). Moreover, trichogynes with branches had male nuclei switching direction at T-junctions, illustrated in Movie S8 (The RFP male nuclei are coloured in yellow in this movie), where the first male nucleus initially migrated right but then reversed and migrated left toward its intended pathway to the protoperithecium. This direction switch of male nuclei at T-junctions has been observed several times in independent experiments (data not shown). These observations led us to conclude that male nuclei are endowed with the capacity to change direction in the course of migration and to perhaps follow an yet unknown signal toward the protoperithecium.

An interesting feature of trichogynes is their ability to fuse with several conidia and as a consequence, to allow the discharge of nuclei belonging to different conidia (Figure 6, Movie S6). It is interesting to note that nuclei from conidium A stopped migration, especially nucleus 1, remaining immobile for 23 min (from t = 1 h 25 min 58 sec to t = 1h 52 min 59 sec) while the nuclei from conidium B were released into the tricogyne. We have measured the average speed of nuclei 1 & 2 before the pause and compared it to their average speed all along the whole trichogyne. Before the pause and across the first 420 µm of migration, the speed of nuclei 1 and 2 were 10.9 and 12.3 µm / min, respectively, compared to 6.9 and 8.0 µm / min respectively, when considering the entire migration of both nuclei. This indicated that the speed of male nuclei is not constant during their migration all along the trichogyne and that migration of some male nuclei may interfere with the migration of other nuclei within the same trichogyne.

## Discussion

Sexual reproduction in fungi and fruiting body development have attracted interest of researchers for centuries (Ainsworth, 1976). Imaging these structures under non-live-cell imaging conditions, such as electronic microscopy, have highlighted the extraordinary complexity of fruiting bodies (Harris et al., 1975; Lord and Read, 2011; Mai, 1976; Read, 1983). However, lack of live-cell biology has been an obstacle to fully understand sexual reproduction from the initial plasmogamy of sexual cells to the final production of sexual spores. In previous *Neurospora* studies, following the work of B. O Dodge who had identified the sexual reproductive organs in the *Neurospora* genus, Backus in 1939 observed the fertilisation of trichogynes by conidia and described it through incredibly precise drawings (Backus, 1939; Dodge, 1927). In particular, Backus observed that trichogynes could be branched and were able to bind to and coil around (presumably fuse with) several conidia of the opposite mating type (Backus, 1939). Almost a century after these pioneer studies, using modern live-cell imaging technologies, we are able to image live conidial fertilisation. Through live-cell imaging, we visualised the stages of *N. crassa* ontogeny for the very first time, encompassing the migration of male nuclei along the entire trichogyne length from their parental conidia to the ascogonium at the core of protoperithecia. Our experimental system contained male and the female partners bearing differentially H1-sGFP or H1-RFP tagged nuclei. To our great surprise, both these reporter proteins (H1-sGFP and H1-RFP) did not mix once nuclei bearing each of these transgenes were together within the same trichogyne cytoplasm. This property makes our experimental model well-suited to describe and demonstrate the striking opposite behaviour of male and female nuclei within trichogyne.

During trichogyne growth, resident female nuclei undergo anterograde movement at a comparable speed to the trichogyne. Whether this movement is supported by cytoplasmic flow as in vegetative hyphae and whether female nuclei divide during trichogyne growth remain to be addressed (Ramos Garcia 2009). Upon cessation of trichogyne growth at the male conidium, female nuclei occupy the entire length of the trichogyne and are clustered regularly, especially at the trichogyne tip. After contact between the trichogyne and the conidium, the female nuclei then only show oscillatory movements. In a strikingly concerted manner, movements of female nuclei stop right before entry of the first male nucleus into the trichogyne and eventually resume showing female nuclei migrating backward towards the protoperithecium. Identifying what controls female nuclei movements during fertilisation will be a great challenge. First clues are given by the observation of female nuclei shape during the different steps described above. Indeed, female nuclei shape changes from oval or pear-shape before entry of male nuclei to round during the immobilisation phase and then oval- or pear-shaped when they move again. These changes may be reminiscent of modifications in the interaction between these nuclei and the cytoskeleton. In filamentous fungi, the microtubule (MT) cytoskeleton as well as its associated molecular motors, *i*.*e* the kinesins and the dynein-dynactin complex, have been involved in finely tuned movements such as shape changes, oscillatory movements, nuclear division and nuclei distribution in hyphae. In particular, both changes of shape as well as oscillatory movements are affected when cells are treated with MT-depolymerizing drugs as well as in the *ropy* mutants altered for the dynein-dynactin complex (Freitag 2004, Lang 2010, Mourino-Perez 2017, Roca 2010, Gladfelter 2009). Moreover, nuclei in the *ropy* mutants as well as in the *kin-1* mutant (altered for the major molecular motor Kinesin-1) have a rounded shape in contrast to the oval- or pear-shape of nuclei in wild type cells. Therefore, it is likely that the oscillatory movement of female nuclei within the trichogyne might involve MT cytoskeleton and the kinesin/dynein molecular machinery. We speculate that change of shape of female nuclei together with their immobilisation may be reminiscent of momentary loss-of-interaction of these nuclei with the MT cytoskeleton machinery during the time male nuclei migrate. In addition, we have observed that the immobilisation of female nuclei may not be trichogyne-autonomous since it was detected in surrounding hyphae. Altogether, these data led us to hypothesize that a diffusible signal that triggers immobilisation of female nuclei is produced at onset of male nuclei entry into the trichogyne.

Contrary to the resident female nuclei, male nuclei have to migrate up the trichogyne in order to reach the ascogonium in the core of the protoperithecium. Here, this assumption has been proven correct via the tracking of male nuclei all along their migration across the trichogyne until the core of the protoperithecium, presumably the ascogonium. We observed that several male nuclei could enter a single trichogyne, raising the question of the poly-fertilisation of protoperithecia. The possibility for more than one male nucleus to fertilize a single protoperithecium has genetically been addressed in *N. crassa* (Nakamura and Egashira, 1961; Sansome, 1949; Weijer and Dowding, 1960). Sansome was the first to demonstrate that ascospores from a single perithecium could originate from more than one pair of parental nuclei. In 1960, Weijer showed that 23% of the perithecia contained rosettes initiated by more than one male nucleus when heterokaryotic male parent were used. However, Nakamura *et al*. found that only 2% (60/2770) perithecia display mosaic rosettes accounting for fertilisation by two or more different male nuclei when a mix of genetically different conidia is used. This shows that using this more “natural” method (very similar to our fertilisation conditions), fertilisation of a single protoperithecium by more than one male nucleus is a rare event. Nonetheless, we show that several male nuclei can enter a single trichogyne, and that they can originate from different conidia, proving that a single protoperithecium can be fertilised by more than one male parent. This is in sharp contrast with the rarity of poly-fertilisation observed by Nakamura *et al*. However, in two cases at least, we have observed that only one male nucleus penetrated the core of the protoperithecium while all the other nuclei within the same trichogyne were blocked. Hence, we speculate that entry of the first male nucleus into the protoperithecium may inhibit entry of the following ones. By comparison with fertilisation in animals, this blockage could be reminiscent of a mechanism avoiding “polyspermy” in the fungus *N. crassa*. The genetic evidences mentioned above show that this blockage may be leaky, leading to poly-fertilisation of protoperithecia in some cases.

Although we ignore how male nuclei find the way to the protoperithecium in a branched trichogyne, we observed male nuclei first making a hesitant turn at T-junctions then reversing to take the other branch, likely the correct path to the protoperithecium. This highlights that male nuclei can rapidly change direction in a trichogyne (moving back-and-forth in a retrograde *vs*. an anterograde movement) and that the migration path is not unique, nor its signalling precisely defined. We speculate that this signalling may regulate the tethering of male nuclei to molecular motors, but also the choice of the appropriate molecular motor to move in the anterograde or the retrograde direction.

The most striking features observed in our attempts to characterise the behaviour of nuclei in the course of fertilisation are indubitably the ones related to male nuclei. Male nuclei undergo a series of morphological changes within the conidium at onset of migration. Since we could not observe when cell fusion between trichogynes and conidia occurs (plasmogamy), how these morphological changes in the nuclei synchronise with plasmogamy remains to be addressed. Next, male nuclei undergo initial stretching upon leaving the conidium. Male nuclei then migrate up the trichogyne repetitively stretching and contracting in an inch-worm-like manner. Generally, the migrating nuclei harbour a compact shape before encountering septa and elongate dramatically while passing through. This stretching can be as extreme as 40 µm length, around 20 times the size of standard nuclei, shedding light to two important features of migrating male nuclei: the extraordinary plasticity of the nuclei and the strong pulling force applied to the nucleus. This plasticity is particularly striking when the rear of the nuclei remains blocked at septa while in the meantime, the front is repetitively pulled leading to repetitive stretching. In the end, like for a rubber band stretched and released at one side, the rear of the nuclei eventually joins the front in a movement exerting the highest speeds ever measured for nuclei in cells (130 µm / min). We propose that the force moving nuclei in the trichogyne is likely a pulling force applied to the front of nuclei and that the rear follows by a spring effect. Since male nuclei can rapidly change direction and move back-and-forth, this imply that the leading ‘front’ of nuclei can become the ‘rear’ and *vice-et-versa* almost instantly.

Regarding the male nuclear velocity, its versatility and the fact that during this impressive mobility, female nuclei remain immobilised, we exclude the cytoplasmic flow to be responsible for male nuclei movements. Hence, we suppose that the movements of male nuclei are undertaken by molecular motors and cytoskeleton (Gladfelter and Berman, 2009). Further studies are required to determine the role of the microtubules, of actin as well as the role of their cognate molecular motors, *i*.*e*. myosin *vs*. dynein/dynactin and kinesin respectively in the movement of male (but also of female nuclei) within the trichogyne during sexual reproduction.

The difference in behaviour of male and female nuclei inside the same fungal cell cytoplasm raises the question of the identity of both types of nuclei. Since heterothallic fungi can only reproduce by sexual reproduction with individuals of opposite mating type, it is assumed that the mating type locus is the primary determinant of the male and the female nuclei identity (Debuchy et al. 2010). In particular, *mat* loci regulate the expression of the pheromones and their cognate GPCRS, which are both involved in the chemotropic interplay between conidia and trichogynes. Basically, binding of the pheromone produced by conidia to its cognate receptor at the trichogyne membrane activates the GPCR signalling cascade responsible for the chemotropic growth, the coiling and ultimately the fusion with conidia of opposite mating type only (Kim and Borkovich, 2004, 2006; Kim et al., 2012; Krystofova and Borkovich, 2006). It will be of great interest to test whether the *mat* locus and the pheromone/GPCR pathway are involved in the control of male and female nuclear movements during sexual reproduction in *N. crassa*.

Through modern live-cell imaging, our work humbly ambitioned to answer questions about nuclei fate in the course of fertilisation as old as the discovery of sex in fungi (Ainsworth, 1976). However, our study broadens the perspectives beyond the simple study of fertilisation by unravelling unexpected behaviour of both male and female nuclei. This live-cell imaging approach to study fertilisation in the model ascomycetes *N. crassa* raises new questions: what are the regulation pathways that control immobilisation of female nuclei? What is the machinery that pulls male nuclei during migration? What signal guides male nuclei during migration? How polyspermy avoidance is set up? How male nuclei entry into protoperithecia triggers perithecial development? What defines nuclei identity? On top of this, the dramatic deformation of male nuclei during migration within the trichogyne and the amenability of model fungi such as *N. crassa* for live-cell-biology and molecular genetics studies make fertilisation in *N. crassa* a new system to study the effect of physical forces and of the deformations they create on nuclei. Understanding how physical forces and constraints applied to nuclei modify the whole nucleus organization and trigger gene expression switches is an emerging area of research (Chalut et al., 2010; Guilluy et al., 2014). The number of human pathologies primarily due to failure in mechanotransduction of signals is constantly increasing (Dauer and Worman, 2009) and recent data show that extreme deformation of nuclei in metastatic cells during migration has a priming effect in aggressiveness of these malignant cells (Golloshi et al., 2019). Therefore, the study of the plasticity of the male nucleus and of its chromatin structure during sexual reproduction in *N. crassa* offers a unique opportunity to unravel how extreme deformations of nuclei can modulate genome structure and gene expression in eukaryotes.

## Materials & Methods

### Strains, culture conditions and production of conidia

*Neurospora crassa* strains used in this the study: wild-type strains *74-OR23-1V A* (FGSC#2489) and *ORS-SL6 a* (FGSC#4200); N2282 *mat A his-3*^*+*^*::Pccg-1-hH1*^*+*^*-sGFP (H1-sGFP* in the manuscript); N2283 *mat a his-3*^*+*^*::Pccg-1-hH1*^*+*^*-sGFP; mat a his-3*^*+*^*::Pccg-1-hH1*^*+*^*-tdimerRed (H1-RFP* in the manuscript). **Construction of the *H1-RFP* strain** (n°263 in Nick Read collection): A plasmid expressing an N-terminal tdimerRed Fluorescent Protein (RFP) fused to histone H1 was constructed by inserting the *N. crassa hH1* gene, amplified from pMF280 (Freitag et al., 2004) with primers hH1SpeF (5’-GCC*ACTAGT*ATGCCTCCCAAGAAGACCGAG-3’) and hH1XbaR (5’-GCC*TCTAGA*TTGCCTTCTCGGCAGCG-3’) into pMF331 (Freitag and Selker, 2005) digested with *Spe*I and *Xba*I. The resulting plasmid, pMF360, was transformed into N623 (*mat A his-3*; (Freitag et al., 2004)) as previously described (Margolin et al., 1997). Positive transformants were selected on minimal medium and screened for expression of tdimerRed-H1 (H1-RFP in the manuscript) under a dissecting fluorescence microscope as previously described (Freitag and Selker, 2005). A transformant (NMF138) was crossed with N3011 (*mata his-2; mus-51::bar+*) to generate homokaryotic progeny. One *mat a* progeny was called strain “263” in the Nick Read collection.

Strains were maintained and grown on solid Vogel’s minimal medium with 2% sucrose at 27°C. Conidia were harvested from 4-to 5-day-old cultures grown at 25°C on Vogel’s medium with constant light, filtered with Miracloth (https://merkmillipore.com), washed in TS (0,05% tween 80/9%NaCl/sterilized water), and stored in TS at 4°C up to 7 days.

### Fertilisation procedure

Petri plates containing 2% agar (tap water) solid medium were prepared and inoculated with the female strain (*mat a* or *A*) and incubated for 5-7 days until protoperithecia had developed. A 2-3 cm^2^ agar plug harbouring protoperithecia was cut and transferred to an empty petri plate. This sample was inoculated with a 1-2.10^6^ conidia.mL^-1^ suspension of the “male” partner of opposite mating type. After 5-6 h incubation at 25 °C in a moisture chamber, the sample was mounted inverted onto a droplet of Vogel’s medium in a 2-well µ-Slide ibidiTreat chamber (https://ibidi.com/) and inspected for fertilisation. Alternatively, after the 5-6h incubation at 25°C, the sample could be kept at 4°C overnight and mounted the day after in the microscopy chamber. In both cases, migrating male nuclei were observed after of 3-5h further incubation at room temperature.

### Low-temperature scanning electron microscopy (LTSEM)

For Figure 1A and 1C, LSA plates overlaid with sterile cellophane (525 gauge uncoated Rayophane, A.A. Packaging, Preston, UK) were used for sample preparation. Figure 1A: three day-old culture of wild type *mat A N. crassa* strain (74*A*). Figure 1C, 7 day-old wild type (74a) *N. crassa* strain inoculated with wild type *mat A male* conidia (74A) (14h incubation after inoculation). Cellophane rectangles carrying the specimen were cut out and attached to the surface of a cryospecimen carrier (Gatan, Oxford, United Kingdom) with a thin layer of Tissue-Tek OCT compound (Sakura Finetek, Torrance, Calif.) as an adhesive and cryofixed by plunging into subcooled liquid nitrogen. The specimen carrier was transferred under low vacuum to the cold stage of a Gatan Alto 2500 cryopreparation system at -180°C and then to the cold stage of a S4700II field emission scanning electron microscope (Hitachi, Wokingham, United Kingdom), where it was warmed to -80°C under continuous visual observation until any surface ice contamination had been removed by sublimation. The specimen was then re-cooled to below -120°C, before being returned to the specimen stage of the Gatan Alto 2500 cryo preparation system at -180°C, where it was coated with about 10 nm of 60:40 gold-palladium alloy (Testbourne Ltd., Basingstoke, United Kingdom) in an argon gas atmosphere. The coated specimen was returned to the cold stage of the SEM and examined at a temperature of <-160°C, a beam accelerating voltage of 2 kV, and a beam current of 10 µA, at working distances of from 12 to 15 mm. Digital images were captured at a resolution of 2,560 by 1,920 pixels by 8-bits using the signal from the lower secondary electron detector and saved in TIFF format.

## Live-cell imaging

### Confocal imaging

fertilisation events were imaged in inverted agar samples on a Leica SP8x confocal microscope equipped with argon and supercontinuum white light laser (WLL) excitation sources. The excitation wavelengths were 488 nm (SGFP;argon laser) and 555nm (RFP;WLL); the emission was collected on HyD detectors for sGFP (standard-496-550nm) and tdimerRed (standard-600-721nm), while a transmitted light PMT was used for brightfield collection. **For Figure 2, 6, 7, Movies S1, S2, S6 and S7**, acquisition settings were as follow: the objective HCX PL APO CS 10x/0.40 dry; 8 stack intervals of 4.3 µm; frame rate 2 min^-1^ 27.4 sec^-1^. **For Figure 3, 5 Movies S3a & b, S5, S8 and S9**, objective HCX PL APO UVIS CS2 63x/1.20 water; Movie **S3a**, 31 stack intervals of 0.7 µm; frame rate 1 min^-1^; Movie **S3b**, 18 stack intervals of 1 µm; frame rate 9,9 sec^-1^. For **Figure 5** Movie **S5**, 16 stack intervals of 0.7 µm; frame rate 4.5 sec^-1^. For Movie **S9**, 28 stacks intervals of 1 µm; frame rate 5.4 sec^-1^. For female nuclei speed measurements, nuclei were segmented into spots with the IMARIS software (Oxford Instruments, UK) and the track speeds for each spot were extracted. **Statistical analysis:** A two-way analysis of variance (ANOVA) with fixed effects for target site (hyphae vs. trichogyne), time of observation (before vs. after nuclei male entry), and their interaction was carried out on all available data. Multiple observations from identical loci were pooled using the arithmetic mean, and data were log-transformed to mitigate the right tailed distribution and to stabilize the variance. Post-hoc tests were corrected for multiple comparison using Holm stepwise procedure to ensure adequate family-wise error rate. A type I error rate of 0.05 was considered for all statistical analyses. Except for Figure 1, 4 and Movie S4, all the pictures and movies shown have been generated with IMARIS as Z-projections with maximum intensity. Figure 2, 4, 5 and 6 were done using the function “montage” of the FIJI software (https://imagej.net/Fiji) (Schindelin et al., 2012). **For Figure 4 Movie S4**, widefield imaging was performed on a temperature-controlled motorised Nikon Te2000 equipped with a Nikon PlanApo VC water immersion 60x/1.2NA objective lens. Fluorescence images were captured on a Hamamatsu Orca-ER (Hamatsu Photonics UK Ltd, UK) CCD camera under pE-1 LED illumination (CoolLED, UK) exciting at 470 and 550 nm for sGFP and tdimerRed, respectively, using a Semrock GFP/DsRes-2x-A-NTE filter cube (Chroma) and MetaMorph software (Molecular Devices). 4D time-lapse stacks were acquired with 3 slices at a stack interval of 1µm with a frame rate of 29.6 sec^-1^

## Supporting information

chemotropic trichogyne growth

male nuclei migration into the trichogyne

female nuclei before male nuclei entry

female nuclei after male nuclei entry

male nucleus morphology changes within the conidium

inch-worm like movement of male nuclei

fertilisation of the protoperithecium by male nuclei-1

fertilisation of the protoperithecium by male nuclei-2

male nuclei-direction swich at T-junction

back and forth movement of male nuclei

## Authors’s contribution

SB: experimentation, data analysis & manuscript writing; HCK: experimentation; CEJ: experimentation; DT: manuscript writing; NR: Project leader & manuscript writing.

## Acknowledgments

We dedicate this article to the memory of our beloved dear friend, Professor Nick Read. All imaging and analysis was performed at the Institute of Cell Biology of the University of Edinburgh and at the Phenotyping Centre At Manchester (PCAM) of the University of Manchester. SB is Assistant Professor employed by the Université de Paris and has received a grant from the Université de Paris for his sabbatical at the MFIG where he performed the experiments. This study was supported by IdEx Université de Paris : ANR-18-IDEX-0001; HCK was PhD student and was funded by Overseas Research Studentship. We would like to thank Kathryn Lord for the permission to use the picture of the young protoperithecium in Figure 1A; Kathryn Lord & Alexander Lichius for their help in finding the origin of the H1-RFP strain used in this paper and Michael Freitag for providing us with the details of the H1-RFP construction; Christophe Lalanne for the statistical analyses.

